# Charge reversal at the Lhcb2 N-terminus impairs phosphorylation and PSI–LHCII complex formation

**DOI:** 10.64898/2025.12.25.696481

**Authors:** Akanksha Srivastava, Christo Schiphorst, Jarne Berentsen, Dana Verhoeven, Jan van Leeuwen, Fiamma Longoni, Francesco Saccon, Emilie Wientjes

## Abstract

State transitions balance excitation-energy distribution between Photosystem I and Photosystem II in higher plants. Stn7-mediated phosphorylation of the N-terminus of the light-harvesting complex II protein Lhcb2 plays a central role in photosynthetic state transitions. However, it remains unclear how the intrinsic charge of this region, independent of its phosphorylation status, influences state transitions and thylakoid membrane organization. Here, we introduced specific charge-altering mutations in the Lhcb2 N-terminus of *Arabidopsis thaliana* in the *lhcb2* knock-out background and analyzed their effects on LHCII phosphorylation, state transition dynamics, PSI–LHCII complex formation, and thylakoid ultrastructure. Substitution of a conserved positively charged arginine with a negatively charged glutamate (R2E) markedly reduced Lhcb1 and Lhcb2 phosphorylation and state transition efficiency, and abolished PSI–LHCII complex formation. In contrast, introducing a negative charge at a downstream position (Q9E) had no detectable effects. Electron microscopy revealed no significant changes in thylakoid organization in either mutant compared to WT Lhcb2 plants. Despite strongly reduced Lhcb1 and Lhcb2 phosphorylation in the R2E mutant, residual state transitions persisted, potentially mediated by Stn7-dependent phosphorylation of other target proteins. Together, these results provide insight into the role of N-terminal LHCII electrostatics in state transitions and thylakoid membrane organization in plants.

## Introduction

Oxygenic photosynthesis is driven by two photosystems, photosystem II (PSII) and photosystem I (PSI), which work in series to convert light energy into chemical energy. The photosystems are associated with specific light-harvesting complexes (LHCs) that increase the absorption cross-section and funnel excitation energy to the reaction centers. In higher plants, Lhca1-Lhca4 are specifically associated with PSI and Lhcb1-6 with PSII (Jansson et al. 1997). The major trimeric light-harvesting complex (LHCII), composed of Lhcb1-3, forms different types of LHCII trimers. Two LHCII trimers, called S-trimer (strongly bound) and M-trimer (moderately bound), are part of the PSII supercomplex and permanently associated with this photosystem (Boekema et al. 1999). In contract, “additional” LHCII complexes are more loosely bound and can serve both photosystems. A fraction of this LHCII pool is thought to be a permanent antenna for PSI, transferring energy to Lhca1-Lhca4 (Benson et al. 2015; Bressan et al. 2018; Bos et al. 2019; Schiphorst et al. 2022). Another fraction of the LHCII pool can move between PSI and PSII, balancing the excitation pressure on the two photosystems in a process called state transitions (Allen 1992; Rochaix 2014).

State transitions are regulated by the reversible phosphorylation of LHCII in response to changes in the redox state of the plastoquinone (PQ) pool. When PSII is overexcited, the PQ pool becomes reduced, activating the LHCII kinase STN7. Conversely, when the PQ pool is oxidized, LHCII is dephosphorylated by the constitutively active phosphatase TAP38/PPH1 (Bellafiore et al. 2005; Pribil et al. 2010; Shapiguzov et al. 2010). STN7 phosphorylates the closely related Lhcb1 and Lhcb2 proteins at the third threonine (T3) of their mature N-terminal sequences. STN7 recognizes the conserved N-terminal motif basic/basic/threonine/variable/basic (B/B/T/X/B) of Lhcb1 and Lhcb2. This motif forms a positively charged region that fits into a negatively charged cleft of STN7 (Liu et al. 2016; Crepin and Caffarri 2018). The motif is especially well conserved in Lhcb2, with the sequence RRTV(K/R) in most vascular plants (Leoni et al. 2013; Crepin and Caffarri 2015; Longoni et al. 2015). Lhcb3 lacks this N-terminal threonine and cannot be reversibly phosphorylated by Stn7 (Jansson 1994; Crepin and Caffarri 2018). Protein phosphorylation adds about 2 negative charges at neutral pH and changes the chemical nature of the protein surface, allowing altered protein-protein interactions (Hunter 2012).

Although the LHCII S-trimer and M-trimer of the PSII supercomplex can be phosphorylated, they are stably associated with PSII. Only a specific pool of mobile LHCII trimers, composed of two Lhcb1 and one phosphorylated Lhcb2, associates with PSI upon phosphorylation of Lhcb2 (Galka et al. 2012; Crepin and Caffarri 2015; Longoni et al. 2015). The structure of the PSI–LHCII supercomplex, resolved by electron microscopy, reveals specific interactions between phosphorylated Lhcb2 (P-Lhcb2) and PSI subunits, supporting the molecular recognition model of state transitions (Allen and Forsberg 2001; Pan et al. 2018; Wu et al. 2023). This model proposes that phosphorylation enhances the affinity of LHCII for PSI while decreasing its affinity for PSII (Allen and Forsberg 2001). In the absence of Lhcb2, or when its phosphorylation site is mutated, the PSI–LHCII supercomplex cannot form. However, such mutants still retain approximately 40% of their state transition capacity (Vayghan 2002; Pietrzykowska et al. 2014; Cutolo et al. 2023), suggesting that alternative mechanisms or partial compensation by other LHCII components, may contribute to residual activity.

The photosystems are embedded in the thylakoid membrane. The thylakoid is arranged into a three-dimensional network consisting of stacked grana and unstacked stroma lamellae. PSII and LHCII are predominantly located in the stacked grana regions, while PSI is enriched in the unstacked regions. The stacking of grana membranes depends on attractive van der Waals forces between the flat stromal surfaces of PSII-LHCII of neighbouring membranes. These attractive forces have to overcome the repulsive electrostatic forces arising from the net negative charge of the thylakoid membrane. This is only possible when positively charged ions, *in vivo* Mg^2+^, screen the negative charges and reduce the repulsive forces. Depletion of Mg^2+^ results in grana unstacking, which can be reversed by reintroducing positive ions. PSI is excluded from these stacked domains, due to its bulky stromal extensions, and instead segregates into the unstacked stromal lamellae (Staehelin 1976; Barber et al. 1980; Chow and Barber 1980; Barber 1982; Day et al. 1984; Kirchhoff et al. 2007; Kiss et al. 2008; Puthiyaveetil et al. 2017).

Several lines of evidence suggest that LHCII is specifically important for grana stacking, with a particular role for the N-terminus. First, reducing the amount of LHCII reduces the level of grana stacking (Bassi et al. 1985; Pietrzykowska et al. 2014; Sattari Vayghan et al. 2022). Second, according to the “Velcro” model, grana stacking is mediated by electrostatic interactions between the positively charged N-termini of LHCII (especially Lhcb1 and Lhcb2) and negatively charged stromal LHCII surfaces on the adjacent membranes (Standfuss et al. 2005; Daum et al. 2010). Furthermore, cation-mediated salt-bridges between negatively charged LHCII residues in opposite membranes can add to the stacking (Wan et al. 2014). Third, when the N-terminus of LHCII is removed in Mg^2+^ depleted destacked thylakoids, then the grana do not restack after re-introduction of Mg^2+^ (Carter and Staehelin 1980).

Reversible LHCII phosphorylation not only regulate state transitions, but also modulates the thylakoid architecture (Kyle et al. 1983; Chuartzman et al. 2008; Wood et al. 2018; Wood et al. 2019). Mutant *Arabidopsis thaliana* (Arabidopsis) plants, in which LHCII is always phosphorylated, have smaller grana compared to WT, while the opposite is true for mutants that lack the LHCII kinase STN7 (Fristedt et al. 2009; Armbruster et al. 2013; Iwai et al. 2018). Light conditions that enhance LHCII phosphorylation are correlated with fewer membrane layers per granum and reduced grana diameter. These adjustments of the thylakoid structure are thought to tune photosynthetic light-harvesting (Wood et al. 2018; Wood et al. 2019; Garty et al. 2024). Experiments with Arabidopsis mutants that lack the LHCII docking site on PSI indicate that LHCII phosphorylation is required for these changes, but not its association with PSI (Wood et al. 2019). In conclusion, LHCII is centrally involved in organizing and dynamically remodeling the thylakoid membrane system in response to changing light conditions.

Thus far, the role of N-terminal LHCII charges in thylakoid structure and organization has been investigated through modulation of its phosphorylation state. As such, it remains unknown whether the net charge of the N-terminal region, irrespective of phosphorylation, can influence LHCII mobility and grana stacking. However, the recent CRISPR/Cas generated mutant lines that specifically lack Lhcb1 or Lhcb2 (Guardini et al. 2021; Sattari Vayghan et al. 2022; Cutolo et al. 2023), allow alteration of specific residues, and as such the charges of LHCII, in a controlled way. In this study, we introduced specific mutations in the N-terminal region of Lhcb2 to alter its charge, while leaving its phosphorylation site intact. We examined the impact of these mutations on LHCII phosphorylation, state transitions, PSI–LHCII complex formation, and thylakoid membrane organization.

We found that the substitution of the neutral glutamine with the negatively charged glutamic acid (Q9E) resulted in normal state transitions and PSI–LHCII complex formation. Instead, mutating the positively charged arginine into the negatively charged glutamic acid (R2E), showed strongly reduced state transitions similar to the *Lhcb2* knock-out (KO), and lacked the PSI–LHCII complex. Lhcb2 phosphorylation in R2E remained low, consistent with disrupted STN7 recognition. Electron microscopy revealed minor changes in grana structure across genotypes, with slightly increased stacking in the R2E mutant compared to wild type (WT), but similar to the *Lhcb2* KO. These results confirm that Lhcb2 phosphorylation is essential for complete state transitions and show that additional negative charge by itself does not alter state transitions. Furthermore, altered charge does not strongly affect the overall thylakoid architecture. Our findings are in agreement with the idea that Lhcb2 phosphorylation is required for state transitions via specific protein interactions, whereas grana stacking is maintained by a broader network of stabilizing forces.

## Materials and Methods

### Plant materials, growth conditions, and construction of N-terminal charged amino acid mutants of Lhcb2

The wild type *Arabidopsis thaliana* (Col-0) and its *Stn7* KO (Bellafiore et al. 2005), and *Lhcb2* KO (Vayghan 2022) mutants were used. The latter was used to generate N-terminal charged amino acid mutants of Lhcb2 in *A. thaliana.* In all complemented lines, Lhcb2 refers specifically to the Lhcb2.2 isoform, which was selected because it is the most abundant Lhcb2 protein, and its N-terminal region is conserved among Arabidopsis Lhcb2 isoforms. A binary vector pCAMBIA-3300, containing the sequence of the Lhcb2.2 gene fused to a C-terminal single HA tag and driven by its putative native promoter, was used for complementation. This transformation generated WT Lhcb2-HA complemented lines that reverted the mutant phenotype (Vayghan 2020). To complement the *Lhcb2 KO* mutant with mutated Lhcb2.2 genes, specific amino acid substitutions were inserted in the coding sequence by overlapping PCR to obtain three Lhcb2.2 mutated genes: In the R2E mutant, the second arginine of the mature Lhcb2 protein was mutated to glutamic acid; in the Q9E mutant, the ninth glutamine of the mature Lhcb2 protein was mutated to glutamic acid; and in the triple charge mutant, the aspartic acid at position 16 and the glutamic acid at positions 26 and 35 were all mutated into arginine. The complemented lines were selected using glyphosate (30 mg ml^-1^) until homozygous transformants were obtained. The Lhcb2 protein level of the homozygous resistant lines was compared to that of the Lhcb2-WT complemented lines by immunoblotting their total protein extracts with both anti-Lhcb2 and anti-HA antibodies. For each complementation, R2E, Q9E and triple charge mutant; the two independent lines with the most comparable Lhcb2 protein expression levels were selected for further analysis.

All plants were grown in a Hettich PRC 1200 WL plant research cabinet with an 8 h light period of 24 °C and a light intensity of 125 μmol photons m^−2^ s^−1^ /16 h dark at 22 °C. The relative humidity was kept constant at 60%. Plants of 4 to 6 weeks of age were used for the experiment.

### Boltz-2 protein structure modelling

To generate a structural model of the Stn7–LHCII complex, we used Boltz-2, a deep-learning diffusion model for biomolecular structure prediction (Passaro et al. 2025). The mature Arabidopsis protein sequences for Stn7, Lhcb1 (two copies), and Lhcb2 (one copy), together with ATP, were used as input. The prediction was run using default Boltz-2 inference settings.

### Pulse-amplitude modulation (PAM) fluorometry

State transition chlorophyll fluorescence measurements were performed on at least 45 min dark-adapted plants (five biological replicates) using a MINI-PAM-II-R fluorometer (Waltz, Germany) according to (Ruban and Johnson 2009). In brief, a low intensity measuring light (ML Int. setting 1) was applied at first, followed by the application of a saturating pulse (SP Int. setting 8 for 0.8 s) at 10 s. At 1 min, both red actinic light (10 μmol photons m^−2^ s^−1^, λ_max_ = 626 nm) and far-red light (FR Int. setting 12, λ_max_ = 743 nm) were switched on. After 5 min, a saturating pulse was given to register F_m_I’ (State I F_m_’). From 7 to 27 min, the far-red light was switched off to induce state II. Shortly after switching the far-red light on again, saturating pulse was given to register F_m_II’ (State II F_m_’). After 20 min of red and far-red light illumination, a saturating pulse was given to measure F_m_I’ again. The extent of PSII antenna adjustment was evaluated as qT = (F_m_I’ - F_m_II’) / F_m_I’, in which the last F_m_I’ was used. The degree of excitation imbalance was calculated as IB = (F_S_I’-F_S_I) / F_o_ (Ruban and Johnson 2009). The restoration of the electron transport balance was evaluated by qS = (F_S_I’-F_S_II’) / (F_S_I’-F_S_II), where F_S_I’ and F_S_II’ are the steady state fluorescence levels in, respectively, State I and State II in absence of far-red light, while F_S_II is the steady state fluorescence in presence of far-red light (Ruban and Johnson 2009). A qS value of 1 indicates that the state I to state II transition completely restored the PSI:PSII excitation balance, while a value of 0 indicates no restoration of the PSI: PSII excitation balance.

### State transition induction and thylakoid isolation

To induce state I, plants were exposed to 10-15 μmol photons m^−2^ s^−1^ of far-red LED light (λ_max_ = 706 nm) and to induced state II, they were exposed to 10-15 μmol photons m^−2^ s^−1^ of blue light (λ_max_ = 470 nm), both for 45 min. Following the state I and state II inductions, the leaves were harvested from the plants, immediately placed on ice, and used to isolate thylakoids. The thylakoids were isolated as per the protocol previously described by Bos et al. (2019) with the addition of 10 mM NaF as phosphatase inhibitor.

### 77K fluorescence emission spectroscopy

Thylakoids from State I and State II plants were frozen in 50% glycerol in B1 buffer (0.4 M sorbitol, 5 mM MgCl_2_, 20 mM Tricine-KOH pH 7.8, 5 mM EDTA, 10 mM NaHCO_3_, and 10 mM NaF) at a concentration of 20 µg chlorophyll ml^-1^ in a glass Pasteur pipette with a diameter of 1 mm. The fluorescence spectra were measured in liquid nitrogen at 77 K on a fluorescence spectrometer (Edinburgh Instruments FS5, Scotland) after excitation at 470 nm (nine technical replicates). The fluorescence was detected in the 600 – 800 nm range with 1 nm steps. To evaluate relative changes in fluorescence from PSI (733 nm), fluorescence spectra were normalized at 685 nm (PSII maximum).

### Polyacrylamide gel electrophoresis (PAGE), phosphoprotein staining, and immunoblotting

The thylakoids (for phosphoprotein staining) or total protein extracts (for immunoblotting) were separated using a 15% standard Laemmli denaturing SDS-PAGE gel (Laemmli 1970).

To stain phosphorylated thylakoid proteins, the SDS-PAGE gels were stained with Pro-Q Diamond stain (Thermo Fisher Scientific) in accordance with the manufacturer’s instructions. The stained gel images were captured using an Ettan DIGE imager (GE Healthcare) with 540 nm excitation and 595 nm emission. The gels were also stained with colloidal Coomassie to detect total proteins, which were then used as a control for Pro-Q Diamond staining.

For immunoblotting, proteins were transferred to a nitrocellulose membrane at 100 V for 1 h. The membrane was stained using Ponceau S to assess loading efficiency and transfer quality and then blocked using phosphate-buffered saline 0.2% (w/v) Tween and 5% milk powder. The membrane was incubated overnight with two primary antibodies, Lhcb2 (AS01003, Agrisera) or HA-tag (3724S, Bioke). A secondary alkaline phosphatase-conjugated antibody was used with the 5-Bromo-4-chloro-3-indoryl Phosphate (BCIP)/ Nitroblue Tetrazolium (NBT) staining solution to form an intense insoluble purple dye at the location of the antibody. Pro-Q staining and western blotting were performed three times using at least two biological replicates.

### Blue native (BN) PAGE

For BN-PAGE, the protocol as described in (Jarvi et al. 2011) was used after solubilization of thylakoids (to a chlorophyll concentration of 0.5 mg ml^-1^) with 1 % final concentration of digitonin. 15 μl of solubilized thylakoid membranes from each sample were loaded on NuPAGE™ 4-12% Bis-Tris Protein Gels (Invitrogen) and electrophoresis was performed at 4 °C in an XCell SureLock Mini-Cell (Invitrogen). Each BN-PAGE was performed twice using two biological replicates.

### Streak

Time-resolved fluorescence spectra were recorded with a streak camera (C10910, Hamamatsu) setup as described in (Stokkum et al. 2008; Farooq et al. 2018; Bos et al. 2023). Prior to the measurements, WT and *Lhcb2* KO plants were dark adapted overnight. The leaves were placed inside a rotating cell, rotating at 1 rpm and moving sideways at 3 rpm. They were excited with a pulsed white light laser (Rock white light laser, Leukos) at a wavelength of 470 ± 10 nm, with 40 μW laser power, a repetition rate of 38 MHz, and a spot size of approximately 200 μm. To close the PSII reaction centres, an additional continuous laser (λ = 532 nm, intensity 0.5 mW, and spot size of approximately 1 mm) was focused on the same spot as the pulsed laser. The chlorophyll fluorescence above 640 nm was spectrally separated with a grating of 150 g/mm (Shamrock, Andor). This creates a streak image in which the fluorescence photon counts are separated by wavelength and arrival time at the detector. First, a streak image was recorded on the dark-adapted leaves. The fluorescence was recorded with 400 ms integration time over 400 integrations, in a time window of approximately 3 ns. Next, the leaves were exposed to 10 μmol m^-2^ s^-1^ blue light (λ_max_ = 488 nm) for 10 min to induce the transition to State II, the image was recorded with the blue light still on. The spectra resulting from these measurements were corrected for background signal and the wavelength-dependent sensitivity of the detector. The data was analysed with R-package TIMP-based software Glotaran (Snellenburg et al. 2012). The analysis resulted in decay-associated spectra (DAS) showing fluorescence spectra associated with specific fluorescence lifetimes. For normalization, the total summed area of the three DASs from 665 nm to 780 nm is set to 1. From the three DASs, two spectra with different fluorescence lifetimes originate from PSII. The final PSII DAS shows the sum of these two DAS and the weighted average fluorescence lifetimes. The relative contribution of the PSI fluorescence was compared between the WT and *Lhcb2 KO* genotypes in dark-adapted state and after acclimation to blue light.

### Transmission electron microscopy

To induce state II, WT plants were exposed to ∼20 μmol photons m^−2^ s^−1^ of blue light (λ_max_ = 470 nm) for roughly 90 min. Single leaves of 5-6 plants of WT, state II-induced WT, R2E, Lhcb2-HA, and *Lhcb2 KO*, were fixed in 2.5% glutaraldehyde + 2% paraformaldehyde in 0.1 M phosphate/citrate buffer (pH 7.2) overnight at RT in the dark. For the first hour of fixation, the state II-induced leaves were kept under the blue light to minimise relaxation. After fixation, the fixative was removed, and the samples were washed three times for 10 min in 0.1 M phosphate/citrate buffer under constant rotation. A ∼1 mm^3^ square section of each leaf was cut and washed three additional times in the phosphate/citrate buffer. Samples were further fixed according to a protocol similar to (Zou et al. 2024). In short, leaf sections were fixed with 1% osmium tetroxide in the phosphate/citrate buffer for 1 h at RT. The sections were then washed 5 times in ultrapure water. The specimens were dehydrated in a graded ethanol series: 10%, 30%, 50% (each 10 min), 70% (overnight), 80%, 90%, 96%, 100% (each 10 min), and finally 100% for 20 min. After dehydration, leaf sections were infiltrated with LR White resin using a graded series (1:2 resin:ethanol, 1:1 resin:ethanol, 2:1 resin:ethanol; each 60 min), followed by 100% resin for 120 min, then overnight, and a final 120 min infiltration in 100% resin. Individual leaf sections were added to gelatin capsules filled with LR White, and the resin was solidified by placing the capsules at 60°C for 22 h. Thin sections were prepared using a Leica ultramicrotome UC7. The sections were stained with 2% uranyl acetate for 10 min, washed in ultrapure water, stained with 3% CO_2_-free lead citrate for 10 min, and washed in ultrapure water. Chloroplasts in the leaf sections were imaged using a JEOL JEM 1400 transmission electron microscope (120 kV).

## Results

To investigate the role of charges in the N-terminus of Lhcb2 on state transitions and grana stacking, we complemented the *Lhcb2* KO line with WT and charge-mutated versions of Lhcb2. An HA tag was added to the C-terminus of Lhcb2 to allow reliable detection of the expressed protein in the complemented line. First, we aimed to mimic Lhcb2 phosphorylation. This can be done by replacing threonine-3 of the mature sequence (also called T40 for the full sequence including the transit peptide) with a negatively charged residue, as previously shown (Vayghan 2022). However, at physiological relevant stromal pH conditions (pH 7.6-8.0 in the light), a phosphate group adds about two negative charges to the subunit. To introduce a larger shift in net charge, we replaced the positively charged arginine preceding threonine-3 with glutamate (R2E), thus changing a conserved positive charge into a negative one while retaining the threonine itself (Fig. 1A, B). In the second mutant, we substituted the neutral residue glutamine at position 9 with the negatively charged glutamate (Q9E). While Arabidopsis and several other plants have a glutamine at this position, *Picea glauca* naturally carries a glutamate (Crepin and Caffarri 2018). We hypothesized that increasing the negative charge of the Lhcb2 N-terminus could alter thylakoid stacking and the rate of state transitions. Finally, we designed a triple mutant (D16R, E26R, E35R) to increase the overall positive charge of the N-terminal region, which we expected to increase grana stacking according to the “Velcro” model. See Figure 1A for the N-terminal sequence of the WT and mutant Lhcb2 and Figure 1B for the charge distribution of the mobile Lhcb1_2_Lhcb2_1_ LHCII complex.

**Fig. 1.**
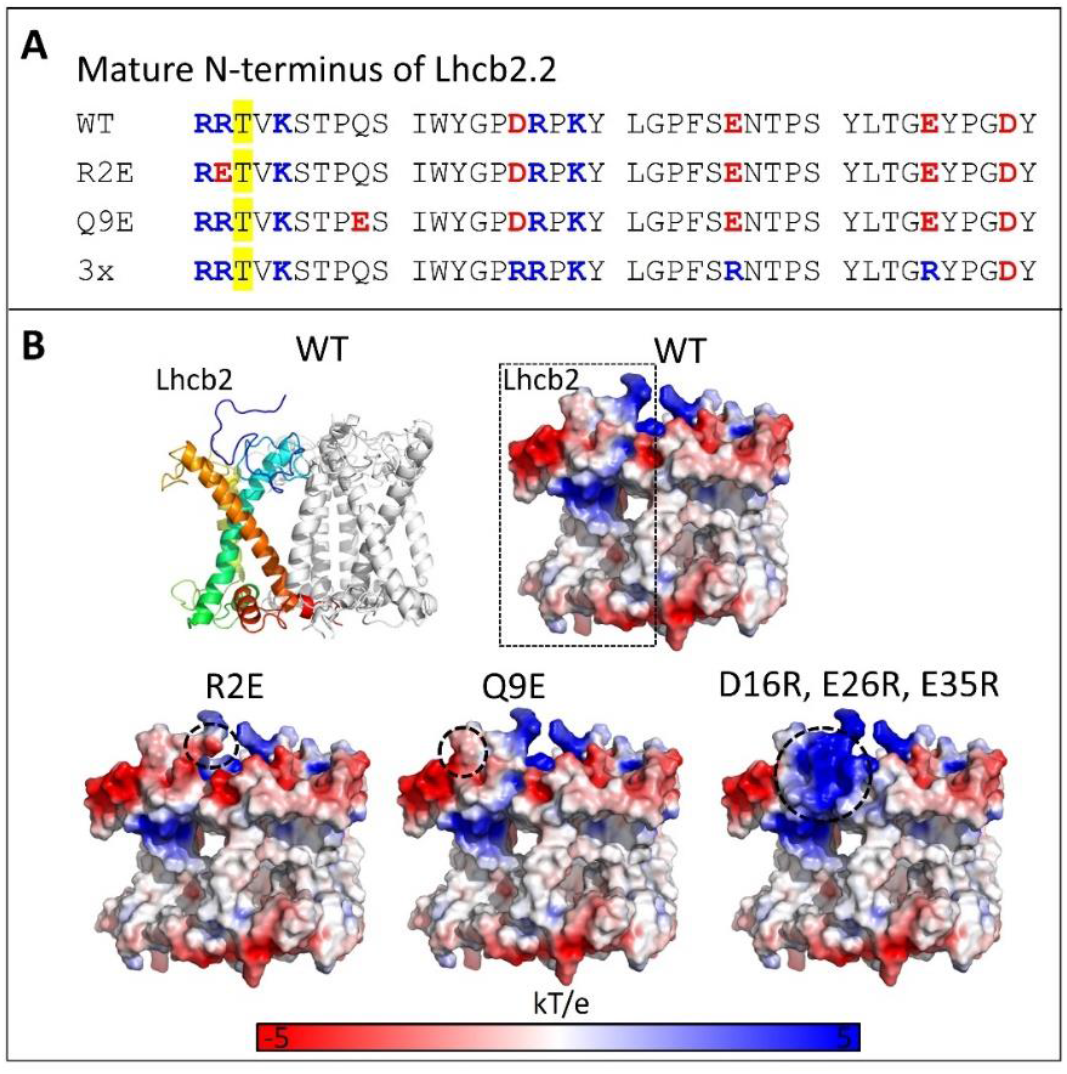
Lhcb2 mutants. A. Mature N-terminal WT and mutant sequences of Lhcb2.2 of Arabidopsis. B. Structure of Lhcb1_2_Lhcb2 LHCII trimer, with Lhcb2 rainbow coloured with a blue N-terminus and the C-terminus in red (left top). The structure was predicted with Boltz-2 (since EM and crystal structures usually lack the complete N-terminus). The predicted model overlaps well with LHCII from the PSI-LHCII complex of Zea mays (Pan et al. 2018), PDB 5ZJI. The other structures show electrostatic surface representations of WT and mutant Lhcb2 within the trimer, where positively charged regions are coloured blue, neutral regions white, and negatively charged regions red, using a potential range of ±3 kT/e. Electrostatic maps were calculated with PDB2PQR (Dolinsky et al. 2004) and visualized in PyMOL. Areas where the charge is altered in the mutants are indicated with a dashed black line.

After generating the HA-tagged charge-mutated Lhcb2 constructs, each variant along with wild type Lhcb2 were introduced into the *Lhcb2* KO background, selected for antibiotic resistance, and subsequently crossed to obtain homozygous lines. Further, immunoblotting with the Lhcb2 antibody, were used to compare the expression levels of Lhcb2 in the mutant plants relative to WT. The expression level of the WT Lhcb2-HA was about 50% of the untagged WT. The R2E-Lhcb2 mutant showed higher Lhcb2 band intensity than WT Lhcb2-HA (comparable to untagged WT), while the Q9E mutant and the triple (D16R, E26R, E35R) mutants showed reduced signals, corresponding to approximately 23–27% and 17–18% of untagged WT levels, respectively (SI Fig. S1). Since the Lhcb2 antibody is specific to the N-terminal region, its affinity for the Lhcb2 mutants may be affected. To obtain a mutation-independent estimate of protein abundance, we took advantage of the altered migration of HA-tagged Lhcb2 relative to Lhcb1 on SDS–PAGE, allowing quantification relative to WT Lhcb2-HA (which expresses at 50% of untagged WT levels; SI Fig. S2). This analysis showed that the R2E and Q9E Lhcb2-HA are expressed at levels similar or slightly higher to WT Lhcb2-HA. No band was detected for the triple mutant, suggesting its expression was limited. Immunoblotting with the anti-HA antibody (SI Fig. S3) confirmed the low expression level of the triple mutant (26–28% of untagged WT), which was therefore not investigated further. The other mutants exhibited R2E Lhcb2-HA expression levels similar to or slightly higher than those of WT Lhcb2-HA (58–77% of untagged WT) and Q9E Lhcb2-HA expression levels similar to or slightly lower than those of WT Lhcb2-HA (35–56% of untagged WT). Although the results obtained using different methods are not fully consistent, they all indicate 50-100% Lhcb2 expression levels for WT Lhcb2-HA, R2E, and Q9E compared to untagged WT, at least in one of the two independently tested lines for both R2E and Q9E.

Next, state transitions were investigated using pulsed-amplitude modulated (PAM) leaf fluorescence (Fig. 2) in dark-adapted WT and mutant plants. The R2E and Q9E mutants were compared with WT, WT Lhcb2-HA, *Stn7* KO (locked in State I) and *Lhcb2* KO plants. First, far-red light was added to weak red light. Far-red light preferentially excites PSI, which oxidizes the PQ pool and results in LHCII dephosphorylation. P-LHCII that was associated with PSI now moves to PSII, resulting in a maximal PSII antenna size, a condition referred to as State I. A saturating pulse at this point provides the maximum PSII fluorescence in the light for State I (F_m_I’). Next, the far-red light was switched off, leaving only the red light on, this reduces the PQ pool and increases the steady state fluorescence level (increase from F_s_I to F_s_I’). Subsequent activation of STN7 results in LHCII phosphorylation, the association of LHCII with PSI, and a reduction of the PSII antenna size (State II). The relocation of LHCII oxidizes the PQ pool, which is reflected by the reduction of the steady state fluorescence level (F_s_II’). The subsequent addition of far-red further oxidizes the PQ pool and immediately decreases the steady state fluorescence (F_s_II). A saturating pulse applied at this moment yields the maximum PSII fluorescence for State II (F_m_II’). The height of the F_m_II’ peak compared to F_m_I’ is directly related to the change in PSII antenna size and thus to the fraction of PSII antenna that moved from PSII to PSI during the State I -> State II transition. Leaving the far-red light on brings the leaf back in State I, allowing to measure F_m_I’ again.

**Fig. 2.**
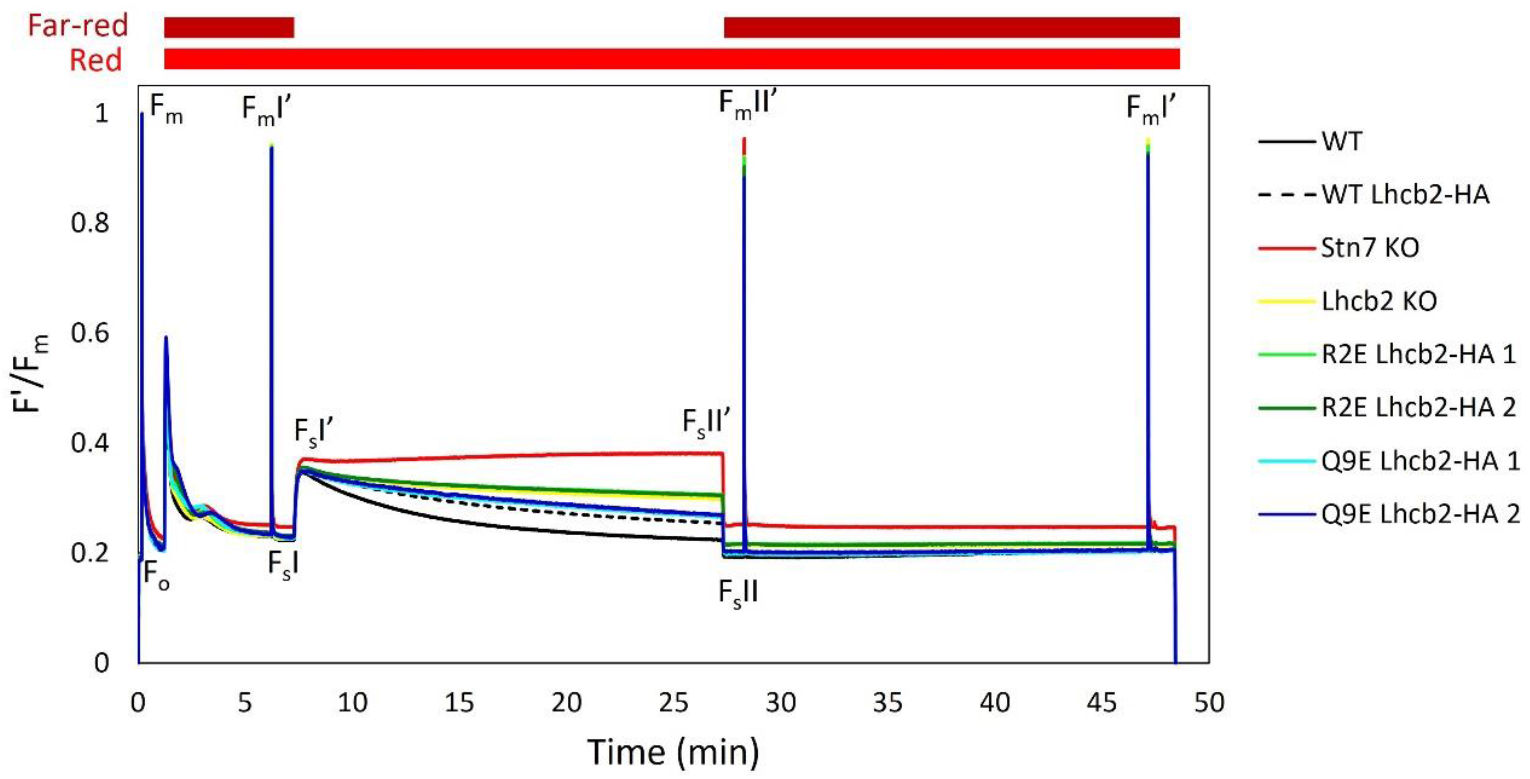
State transitions. Pulse amplitude modulated (PAM) fluorescence traces to compare state transition of WT, WT Lhcb2-HA, Stn7 KO, Lhcb2 KO, R2E Lhcb2-HA (2 independent lines) and Q9E Lhcb2-HA (2 independent lines) plants. The red light (10 μmol photons m^−2^ s^−1^) and far-red light were switched on directly after the F_o_ and F_m_ measurement. The far-red light was switched off from minute 7 to minute 27, as indicated in the figure. The presented fluorescence traces show the average of five biological replicates.

The PSI:PSII excitation imbalance was calculated based on the change in the steady state fluorescence after switching off the far-red light (F_S_I’-F_S_I), relative to the F_o_ level (Ruban and Johnson 2009). When the PSII antenna size is strongly reduced, as in the Lhcb1-deficient line, PSI remains over-excited and the imbalance after changing the light conditions approaches zero (Pietrzykowska et al. 2014). In contrast, all plants tested here showed a similar imbalance (Fig. 3A). Therefore, there is a need for state transitions to restore the excitation balance, e.g. reducing the antenna size of PSII while increasing that of PSI.

**Fig. 3.**
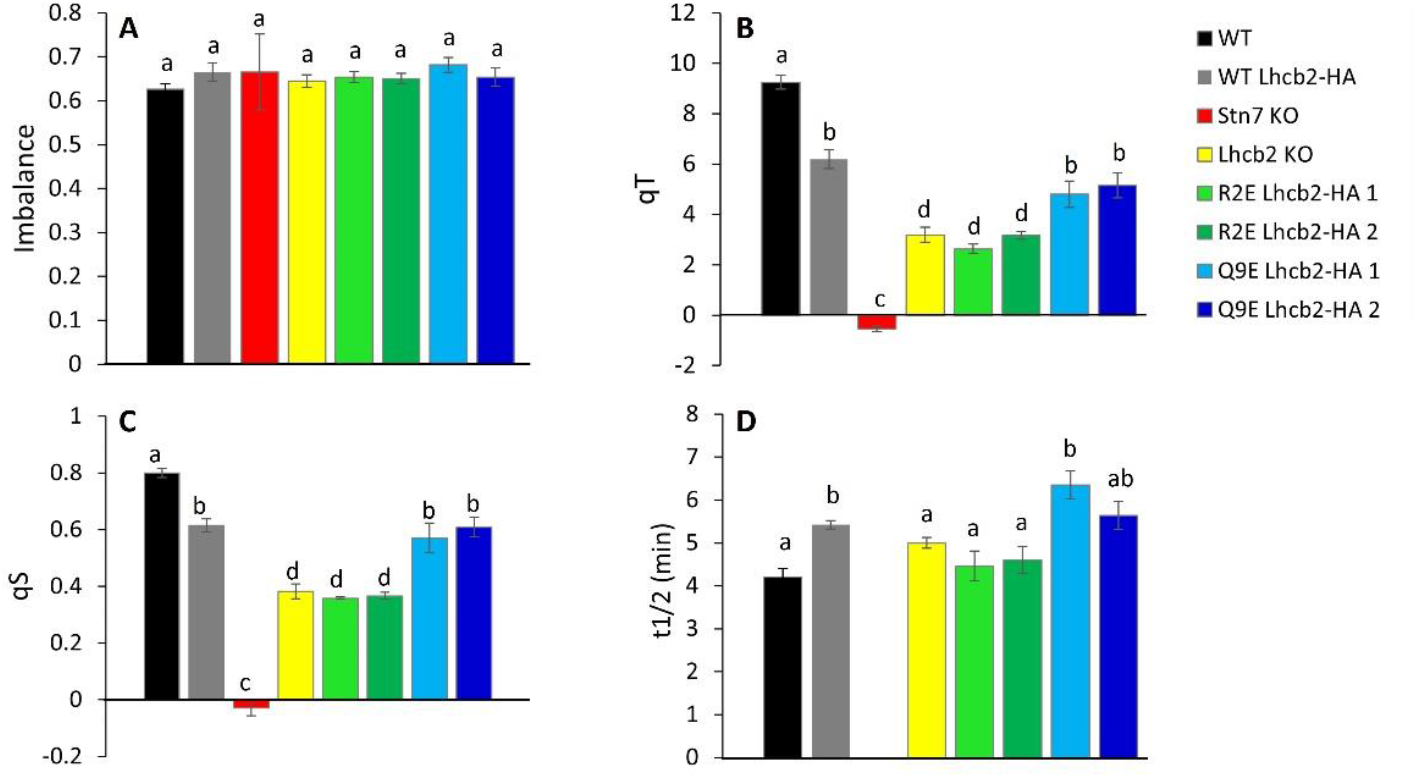
Quantification of state transitions. The imbalance (A), qT value (B), qS value (C) and State I -> State II half-life (D) are given with their SE, calculated from five biological replicates derived from the traces shown in Figure 2. Statistically different values (p < 0.05) are indicated with different letters. Statistical analysis was performed using one-way ANOVA with post-hoc Tukey test.

Next, the change in PSII antenna size during state transitions (qT) was calculated based on the change in F_m_’ value upon the State II to State I transition (Fig. 3B). The WT Lhcb2-HA line had a lower qT than WT plants, which is probably due to the lower Lhcb2 expression level (SI Fig. S1). The *Lhcb2* KO plants showed ∼30% state transitions compared to WT, in agreement with earlier results (Vayghan 2022; Cutolo et al. 2023), although this residual activity is unexpected given that PSI–LHCII complex formation normally requires phosphorylated Lhcb2 (Cutolo et al. 2023). The *Stn7* KO had a qT value close to zero, in line with the absence of the STN7 LHCII kinase (Bellafiore et al. 2005). Among the tested charge mutants, the R2E lines behaved similarly to the *Lhcb2* KO, while the Q9E lines showed a response comparable to WT Lhcb2-HA. Finally, 77K fluorescence emission spectra were used to assess PSI versus PSII antenna distribution in State I and State II. These measurements supported the qT results across all tested plants (SI Fig. S4).

State transitions modulate the relative electron transport through PSII and PSI in response to changes in light quality. The parameter qS quantifies the efficiency of this adjustment, with a value of 1 indicating complete rebalancing of electron flow after a spectral shift, and a value of 0 indicating no adjustment at all, meaning that the photosynthetic apparatus does not respond to the change in the light quality (Ruban and Johnson 2009). In our study, the qS values followed the overall same pattern as the qT values (Fig. 3C). WT plants showed the highest qS, followed by the WT Lhcb2-HA and the Q9E mutant. The *Lhcb2* KO and R2E Lhcb2-HA line show lower values, but still about 2/3th of the WT Lhcb2-HA line and halve of the WT plants, indicating the excitation balance is still partly restored. Instead, no restoration is observed for the *Snt7* KO plants.

Altering the charge of Lhcb2 could influence the rate at which LHCII moves between PSII in State I and PSI in State II. To assess this, we quantified the half-life of the reduction of the steady state fluorescence signal after switching off the far-red light (Fig. 3D, State I -> State II; see SI Fig. S5 for the normalized traces). A shorter half-life corresponds to a faster state transition. Unexpectedly, all lines showed half-lives similar to WT and WT Lhcb2-HA, ranging from 4.2 ± 0.2 min (average ± SE, n=5) in WT to 6.4 ± 0.3 min in one of the Q9E Lhcb2-HA lines. Even in the *Lhcb2 KO*, the half-life (5.0 ± 0.1 min) did not significantly differ from the WT. Likewise, the negative charges introduced in the R2E and Q9E Lhcb2 mutants did not affect the State I to State II transition rate (half-lives of 4.5-4.6 ± 0.3 min for the R2E lines and of 5.6-6.4 ± 0.3 min for the Q9E lines, compared to 5.4 ± 0.1 for the WT Lhcb2-HA line).

To assess the formation of the State II-specific PSI–LHCII supercomplex, we performed digitonin solubilization of thylakoids isolated from leaves adapted to State I (45 min of far-red light) or State II (45 min of blue light), followed by native PAGE (Fig. 4A). The PSI–LHCII complex was detected under State II conditions in WT, WT Lhcb2-HA, and Q9E Lhcb2-HA lines, but was absent in the *Lhcb2* KO, *Stn7* KO, and the R2E Lhcb2-HA lines. The absence of the complex in the *Lhcb2* KO confirms earlier findings (Vayghan 2022; Pietrzykowska et al. 2014; Cutolo et al. 2023), supporting the essential role of Lhcb2 in PSI–LHCII assembly. Likewise, the absence of the supercomplex in the *Stn7* KO reflects the requirement of STN7-dependent Lhcb2 phosphorylation for PSI–LHCII formation (Bellafiore et al. 2005). The inability of the R2E mutant to form the complex may result from impaired phosphorylation of T3 by the STN7 kinase, due to the absence of the highly conserved upstream arginine residue that normally promotes efficient kinase recognition. Alternatively, T3 may still be phosphorylated, but the resulting phosphorylated Lhcb2 (Lhcb2-P) might be unable to support a stable interaction with PSI.

**Fig. 4.**
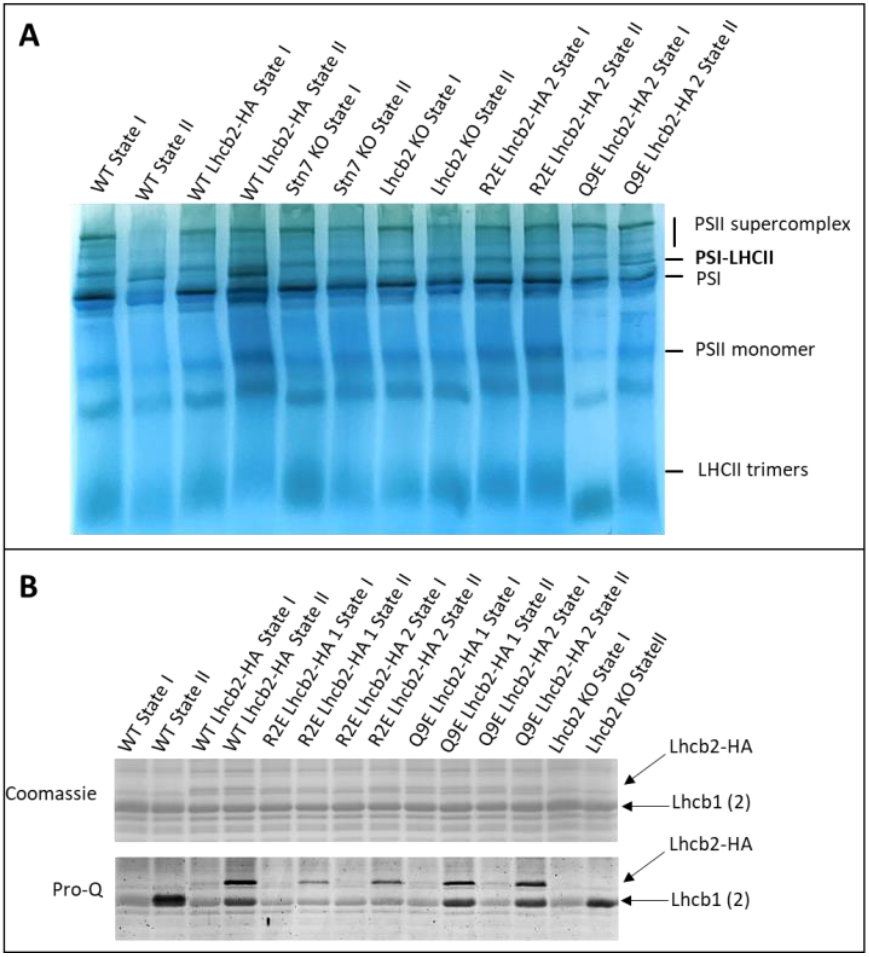
Digitonin native PAGE and Lhcb phosphorylation. A. Native PAGE of digitonin-solubilized thylakoids isolated from leaves under State I (45 min of far-red light) or State II (45 min of blue light) conditions. Samples are as indicated, with two independent lines for each of the R2E and Q9E mutants. The positions of the photosynthetic complexes are indicated. See SI Fig. S6 for the entire gel. B. The same SDS-PAGE gel stained with either Coomassie brilliant blue (top) or Pro-Q phosphoprotein gel stain (bottom). Samples are as indicated, with two independent lines for the R2E and Q9E mutants. The position of Lhcb1, Lhcb2-HA and Lhcb2 of WT band are indicated. See SI Fig. S7 for the entire gels.

To assess the phosphorylation status, we performed Pro-Q phosphoprotein staining. Because Lhcb2-HA migrates differently from Lhcb1 on SDS-PAGE, the two isoforms could be distinguished (Fig. 4B). Under State I conditions, Lhcb1 phosphorylation was weak in all lines, whereas Lhcb2 phosphorylation was nearly undetectable. Under State II conditions, phosphorylation of Lhcb2-HA was strongest in the WT and Q9E lines, whereas the R2E mutant showed a detectable but weaker signal and thus reduced phosphorylation. Lhcb1 was phosphorylated in the WT Lhcb2-HA, Q9E Lhcb2-HA, and *Lhcb2* KO lines. Notably, in the R2E line, Lhcb1 phosphorylation showed little to no difference between State I and State II, with consistently weak signals under both conditions. This suggests that the negative charge introduced in the R2E Lhcb2-HA mutant somehow inhibits the normal induction of Lhcb1 phosphorylation during State II.

To assess whether the R2E mutation in Lhcb2 could directly influence phosphorylation by altering its interaction with the thylakoid kinase Stn7, we generated a structural model of the Stn7–LHCII complex using Boltz-2, based on mature Arabidopsis Stn7, Lhcb1 (2×), and Lhcb2 (1×) sequences and ATP. The model places Stn7 in contact with LHCII via its single transmembrane helix (Fig. 5A), consistent with previous studies indicating that Stn7 contains a single transmembrane helix (Depege et al. 2003; Lemeille et al. 2009).

**Fig. 5.**
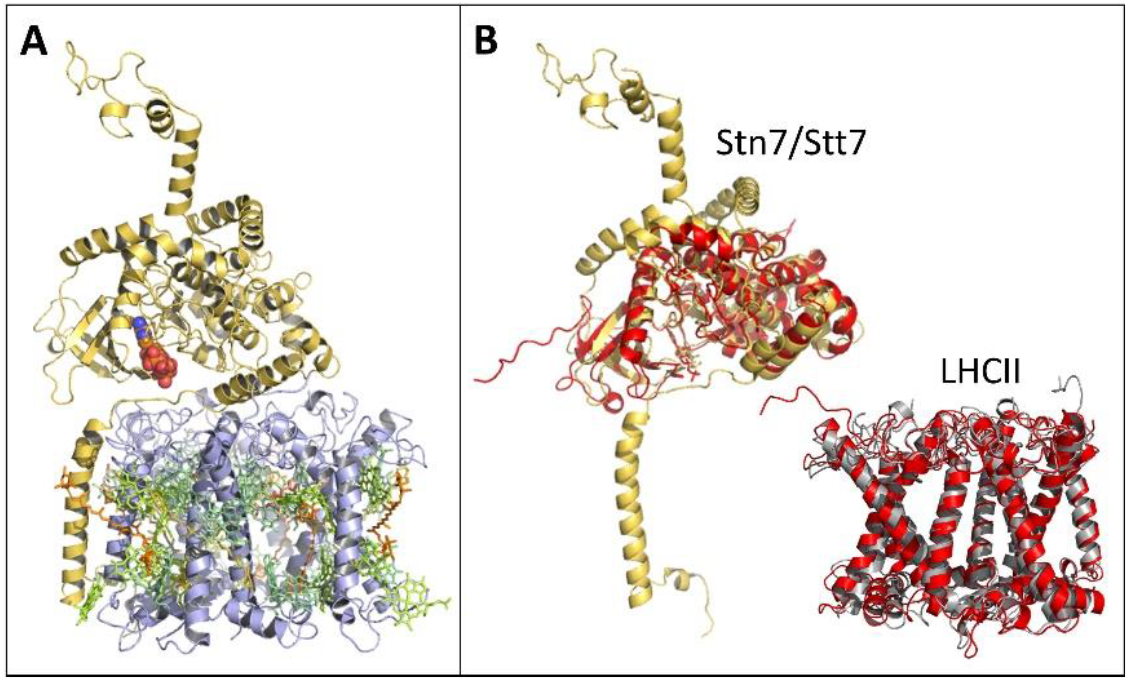
Stn7-LHCII model. A. Boltz-2 model of Stn7-LHCII complex, based on the mature Stn7, Lhcb1, and Lhcb2 Arabidopsis sequences. The pigments are extracted from LHCII of the PSI-LHCII complex from Zea mays (Pan et al. 2018), PDB 5ZJI. Stn7 is displayed in yellow, LHCII in grey, chlorophylls in green, carotenoids in orange and ATP is coloured by element and shown as spheres. B. Alignment of Stn7 model (yellow) with the structure (red) of the catalytic domain of a putative Stt7 kinase homolog from Micromonas algae (Guo et al. 2013) (PDB 4IX6), and alignment of the Lhcb1_2_Lhcb2 trimer model (grey) with LHCII from the PSI-LHCII structure (red).

Furthermore, the catalytic part of Stn7 aligned well with the catalytic domain of a putative Stt7/Stn7 kinase homolog from *Micromonas* algae ((Guo et al. 2013); Fig. 5B), and the modelled Lhcb1_2_Lhcb2 trimer aligned well with LHCII from PSI-LHCII ((Pan et al. 2018); Fig. 5B). We further assessed whether WT and mutant Lhcb2 sequences differed in their interaction with Stn7; however, in neither case did the Lhcb2 N-terminus display stable docking within the Stn7 catalytic site. As such, the model did not allow us to elucidate whether Stn7 substrate recognition was reduced in the mutant. Despite this, our model visualizes the potential interaction between Stn7 and LHCII and provides insight into their relative sizes. The model indicates that the stromal domain of Stn7 extends several nanometres into the aqueous phase, consistent with its localization in the unstacked stroma lamellae (Wunder et al., 2013), where sufficient stromal space is available.

To assess whether the residual state transitions observed in the R2E mutant and *Lhcb2* KO by PAM reflect a State II increase in PSI antenna size or quenching of PSII fluorescence, we performed ultrafast time-resolved chlorophyll fluorescence measurements using a streak camera setup on WT and *Lhcb2* KO leaves. Leaves were adapted to State I (dark) or State II (blue light, 10 μmol m^-2^ s^-1^) conditions and then excited with a pulsed laser that closes PSII reaction centers. This approach enables separation of the rapid PSI fluorescence decay from the slower PSII decay, allowing for analysis of Decay Associated Spectra (DAS). These spectra represent the contribution of individual fluorescence lifetimes at different wavelengths, with characteristic decay times of ∼70 ps for PSI and ∼1.6 ns for closed PSII (Fig. 6A; (Bos et al. 2019)). The relative area under each component reflects its contribution to the total excitation energy.

**Fig. 6.**
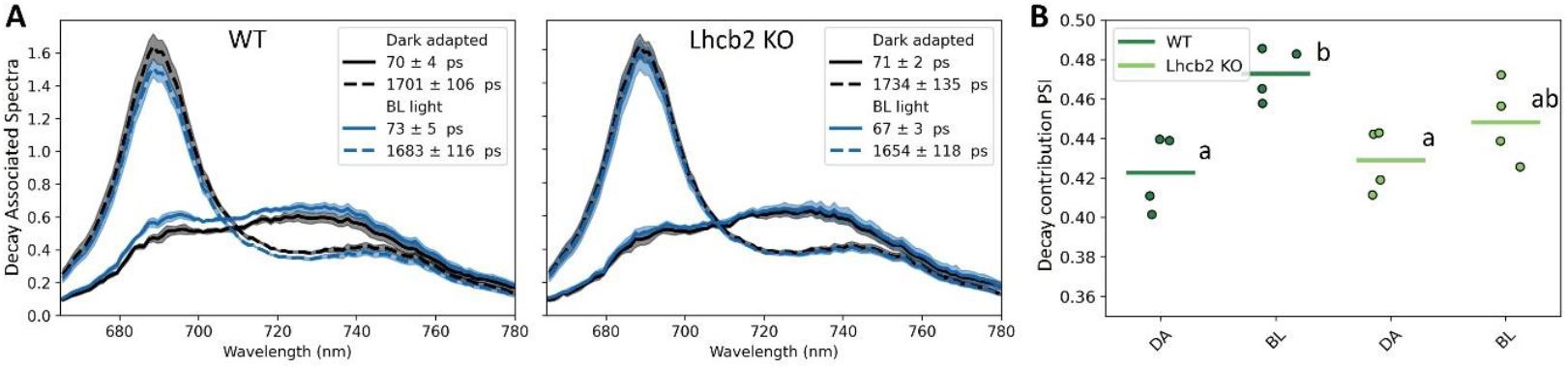
Time-resolved fluorescence analysis. A. Decay associated spectra (DAS) of WT and Lhcb2 KO leaves, measured with excitation wavelength 488 nm (n=4, independent sets of leaves). The leaves were first measured in dark adapted state (State I, grey lines). Then the leaves were adapted to blue light (10 μmol m^-2^ s^-1^) for at least 10 min, and another spectrum was measured (State II, blue lines). The total summed area of the 3 DAS from 665 nm to 780 nm is set to 1. The shaded area shows the SE. B. The contribution of the PSI fluorescence component (∼70 ps) to the total. The letters indicate statistical significance based on a one-way ANOVA followed by pair-wise T-tests (p < 0.05)

In WT leaves, the PSI-associated DAS component increased upon transition from State I to State II, consistent with antenna enlargement of PSI (Fig. 6A, B). In the *Lhcb2* KO, a similar trend was observed, but the increase was not statistically significant, and only minor changes in PSII lifetime were detected under State II conditions relative to State I (SI Fig. S8). Therefore, based on this experiment, it is not clear if increased energy transfer to PSI, quenching of PSII, or both mechanisms are responsible for the remaining state transitions observed with PAM in the *Lhcb2 KO*.

To determine whether altering the N-terminal charge of Lhcb2 affects thylakoid membrane stacking and overall grana architecture, we analyzed chloroplast ultrastructure by transmission electron microscopy (Fig. 7A). This analysis was motivated by models proposing that electrostatic interactions involving the positively charged N-terminus of LHCII contribute to grana stacking, such as the “Velcro” model (Standfuss et al. 2005; Daum et al. 2010). Specifically, we examined the thylakoid organization of the R2E mutant, in which a positive charge is replaced by a negative one, resulting in a net charge of -2, similar to the charge shift induced by T3 phosphorylation. The dark-adapted R2E mutant was compared with the dark-adapted WT (WT_D), WT Lhcb-HA, and *Lhcb2* KO. In addition, WT plants were adapted to weak blue light (∼20 μmol photons m^−2^ s^−1^, λ_max_ = 470 nm, ∼90 min) to induce LHCII phosphorylation (WT_Blue). The number of thylakoid membranes per granum was quantified and expressed in a histogram as percent occurrence (Fig. 7B). As expected, the blue light-adapted WT (State II) had more doublets (grana of two layers) than the dark-adapted WT, in line with previous reports (Pietrzykowska et al. 2014; Garty et al. 2024). In contrast, the R2E mutant showed a WT_D level of doublets, which may reflect either the slightly reduced Lhcb2 abundance in this line (∼77% of WT), causing ambiguity in the interpretation of the results, or the fact that introduction of a negative charge on Lhcb2 alone is insufficient to increase doublet formation. Interestingly, the R2E mutant showed a higher occurrence of >8 thylakoids per granum than the WT or the *Lhcb2* KO, contradicting the hypothesis that the negative charge introduced on Lhcb2 would reduce grana stacking.

**Fig. 7.**
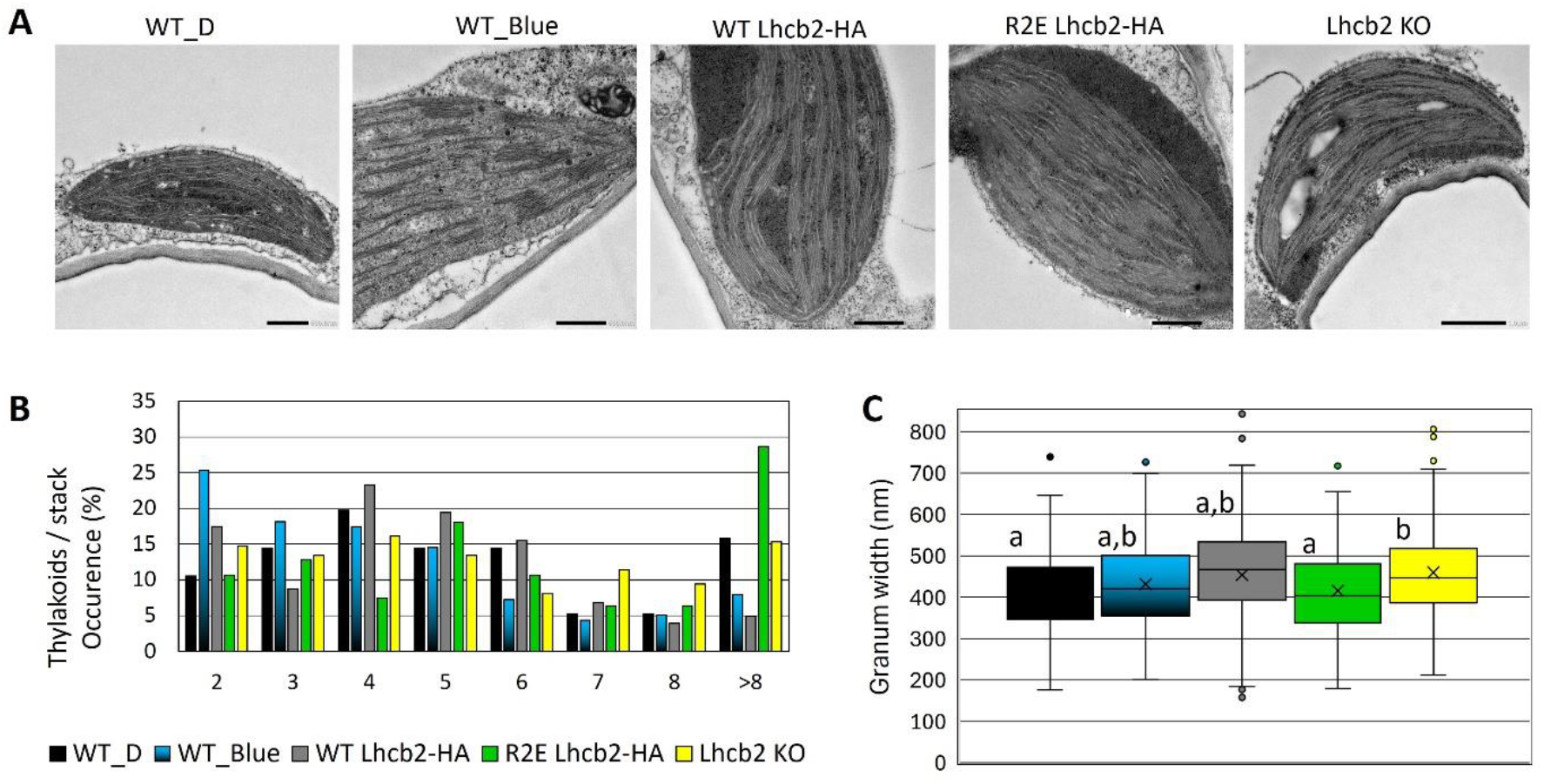
Electron microscopy of the thylakoid membrane. A. Representative EM images of chloroplasts from WT and Lhcb2 mutant leaves: WT dark-adapted (WT_D), WT blue light-adapted (WT_Blue), dark-adapted WT Lhcb2-HA, R2E Lhcb2-HA,and Lhcb2 KO. The scale bar indicates 500 nm. B. Histogram showing the number of thylakoids per granum stack. C. Box-plot of the granum width, 50% of the data falls in the box and 95% of the data falls within the whiskers, outliers are shown as dots. Statistically different values (p < 0.05) are indicated with different letters. Statistical analysis was performed using one-way ANOVA with post-hoc Tukey test. Chloroplasts from at least three plants were analyzed, with 76–152 grana quantified per group.

The granum width was also measured and presented as a box plot (Fig. 7C). The average granum width for WT_D, WT_Blue, WT Lhcb2-HA, R2E Lhcb2-HA, and the *Lhcb2* KO was 412 ± 12 nm (average ± SE), 432 ± 9 nm, 453 ± 13 nm, 415 ± 11 nm, and 460 ± 10 nm, respectively. Unexpectedly, WT plants in State II did not show a reduction in granum width reported previously (Wood et al. 2018; Wood et al. 2019). The only significant differences in grana width were found between WT_D and *Lhcb2* KO, and between R2E Lhcb2_HA and *Lhcb2* KO, with the *Lhcb2* KO showing a slightly larger granum width. These results further indicate that complete loss of Lhcb2 has a more pronounced impact on grana architecture than partial reduction or charge modification of Lhcb2.

## Discussion

To investigate how the N-terminal charge of Lhcb2 affects its phosphorylation, thylakoid organization, and the regulation of state transitions, we introduced specific charge-altering mutations in its conserved N-terminal region. The R2E substitution in Lhcb2, which replaces the conserved N-terminal arginine with a negatively charged glutamate (Fig. 1), markedly reduces but does not abolish Lhcb2 phosphorylation in State II. This suggests that the mutation weakens, rather than completely prevents, recognition by the Stn7 kinase. Despite this partial phosphorylation, no PSI–LHCII complexes were detected on native PAGE, and the qT and qS values were comparable to those of the *Lhcb2* KO line. In contrast, the Q9E substitution, which only adds a negative charge and slightly changes the electrostatics (Fig. 1), did not affect the kinetics of state transitions or the assembly of PSI–LHCII complexes. Therefore, it can be concluded that a moderate change in the N-terminal charge (as in Q9E) does not significantly influence the state transition process (Fig. 2-4).

Surprisingly, in the R2E Lhcb2-HA mutant, phosphorylation of Lhcb1 in State II was also strongly reduced, suggesting a regulatory interplay between the phosphorylation status of Lhcb2 and the activity of Stn7 toward Lhcb1 (Fig. 4B). This observation is in contrast with previous findings for the Lhcb2 T3D phosphomimetic (with one negative charge introduced), which retained normal Lhcb1 phosphorylation levels (Vayghan 2022). While Lhcb1 can be phosphorylated, the LHCII complex associated with PSI in State II consists of one fully phosphorylated Lhcb2 and two unphosphorylated Lhcb1 polypeptides (Crepin and Caffarri 2015; Longoni et al. 2015). Therefore, it is plausible that Stn7 phosphorylates maximum one Lhcb subunit per trimer. In that case, the R2E mutant could mimic the charge of phosphorylated Lhcb2 and thereby hinder Lhcb1 phosphorylation. However, this hypothesis does not fully explain the results, as it fails to account for why LHCII trimers lacking Lhcb2, such as Lhcb1 homotrimers or Lhcb1/Lhcb3 heterotrimers (Jansson et al. 1997; Crepin and Caffarri 2018), are not phosphorylated on Lhcb1.

Both the R2E Lhcb2-HA and *Lhcb2* KO lines lacked detectable PSI–LHCII complexes, as previously shown for the *Lhcb2* KO (Vayghan 2022; Cutolo et al. 2023). Nonetheless, analysis of the PAM traces revealed that these plants still exhibited partial state transitions (Fig. 2), with qT values of approximately 3% (versus 9% in WT and 6% in WT Lhcb2-HA) and qS values of about 0.37 (versus 0.80 for WT and 0.61 in WT Lhcb2-HA). In contrast, in the absence of Stn7, both qT and qS values were close to zero. These residual state transitions in the absence of functional Lhcb2 phosphorylation could potentially be explained by: (i) Lhcb1 phosphorylation-induced quenching of LHCII fluorescence, (ii) excitation-energy spillover from PSII to PSI, (iii) transient associations of phosphorylated Lhcb1-containing trimers with PSI, and/or (iv) phosphorylation of other proteins by Stn7 that affect the PSI:PSII excitation-energy distribution, for example by influencing the thylakoid architecture (Cutolo et al. 2023).

To further explore these mechanisms, we measured time-resolved fluorescence decay in WT and *Lhcb2* KO leaves under State I and State II conditions to distinguish between PSII–LHCII fluorescence quenching under F_m_ conditions and redistribution of excitation energy between PSI and PSII (Fig. 6). However, the small differences observed between the two states in the *Lhcb2* KO were not statistically significant, making it impossible to clearly discriminate between these possibilities with this approach. Nevertheless, only a redistribution of excitation energy between PSI and PSII could explain the PAM data, since the observed changes in fluorescence cannot be attributed solely to quenching processes. The steady-state fluorescence (F_s_′) was 15% lower in State II (F_s_II′) compared to State I (F_s_I′) (Fig. 2). This decrease can be attributed to excitation-energy transfer from LHCII (or PSII) to PSI during the transition to State II, which lowers PSII excitation pressure while enhancing excitation of PSI, a weakly fluorescent photosystem. If, instead, this 15% reduction in fluorescence were due to PSII–LHCII quenching in F_s_′, the F_m_′ quenching should have been even stronger. A quencher causing a 15% reduction in F_s_′ (shorter excited-state lifetime) would induce a proportionally larger reduction in F_m_′, where the excited-state lifetime is longer (see calculation in SI Table S1). Instead, the reduction in F_m_II′ was only 3% relative to F_m_I′. These data therefore indicate that excitation energy redistribution between PSI and PSII still occurs during state transitions even in the absence of Lhcb2 phosphorylation.

In the absence of Lhcb2, the residual state transitions observed by PAM cannot be attributed to Lhcb1 phosphorylation alone. Cutolo et al. (2023) demonstrated that mutation of the Stn7-phosphorylatable threonine in Lhcb1 to valine (which cannot be phosphorylated) does not affect state transitions when WT Lhcb2 is present. This is further supported by the R2E Lhcb2-HA mutant, in which phosphorylation of both Lhcb1 and Lhcb2 in State II was weak, yet the state transition phenotype was indistinguishable from that of the Lhcb2 KO (Fig. 2-4). Future experiments could determine which fraction of Lhcb1 and Lhcb2 proteins remains phosphorylated in State II in the R2E mutant relative to WT, as described by (Crepin and Caffarri 2015; Longoni et al. 2015), and whether the occurrence of thylakoid doublets under State II conditions increases relative to State I, as changes in thylakoid organization have been proposed to regulate excitation-energy distribution and the balance between cyclic and linear electron flow (Wood et al. 2019; Garty et al. 2024). Although Stn7-mediated LHCII phosphorylation is the primary driver of canonical state transitions, our data are consistent with the possibility that Stn7-dependent phosphorylation of other target proteins might contribute to the residual state transitions observed in the Lhcb2 KO and R2E mutants. Notably, NSI-mediated lysine acetylation of chloroplast proteins has recently been shown to be essential for state transitions in general, as *nsi* mutants completely lack PSI–LHCII complex formation and exhibit no detectable state transitions despite normal LHCII phosphorylation (Koskela et al. 2018; Koskela et al. 2020). The presence of residual state transitions in the Lhcb2 KO and R2E mutants therefore indicates that these responses occur in an NSI-competent background but are not directly driven by NSI-dependent acetylation. Instead, NSI function appears to be permissive for excitation-energy redistribution, while the residual state transitions observed here must arise from mechanisms independent of canonical PSI–LHCII complex formation.

Building on the possibility that factors beyond LHCII phosphorylation contribute to residual state transitions, we next asked whether altering the N-terminal charge of Lhcb2 is sufficient to drive the thylakoid ultrastructural changes typically associated with State II. Comparison of the thylakoid organization by EM showed that State II WT plants had more doublets than dark-adapted (State I) plants, supporting previous results (Pietrzykowska et al. 2014; Garty et al. 2024). If this change was only due to the negative charge introduced by Lhcb2 phosphorylation, then a shift towards more doublets would be expected for R2E Lhcb2-HA relative to the WT Lhcb2-HA. However, this difference was not observed. Instead, the R2E mutant exhibited a WT_D-like distribution of doublets (Fig. 7), which may reflect its slightly reduced Lhcb2 abundance (∼77% of WT) and/or indicate that introduction of a negative charge at the Lhcb2 N-terminus alone is insufficient to trigger State II–like thylakoid reorganization.

In conclusion, our study shows that a moderate change in the N-terminal charge (Q9E) does not affect state transitions. In contrast, altering the conserved positively charged residue preceding the Lhcb2 phosphorylation site into a negatively charged residue (R2E) reduces Lhcb2 phosphorylation and, unexpectedly, almost eliminates State II-induced Lhcb1 phosphorylation, thereby significantly impairing state transition efficiency. Even with this low level of LHCII phosphorylation, the residual state transition is comparable to that in the *Lhcb2* KO, in which Lhcb1 is phosphorylated in State II. By contrast, in the absence of Stn7, state transitions are completely lost. Taken together, these findings suggest the involvement of an Stn7-dependent, non-Lhcb protein that is required to achieve WT-level state transitions.

## Acknowledgements

We thank Jelmer Vroom from the Wageningen Electron Microscopy Centre for making the EM images and giving advice during the EM sample preparation.

## Funding

This work was supported by the Dutch Organization for Scientific Research (NWO) via an NWO-M grant (OCENW.M.21.255) and Vidi grant (VI. Vidi 192.042) to E.W and by the Experimental Plant Sciences graduate school through a PhD grant to J.B..

## Declaration of generative AI in the writing process

During the preparation of this work the authors have used ChatGPT to improve language and readability. After using this tool, the authors reviewed and edited the content as needed and take full responsibility for the content of the publication.

## Notes

### Competing Interest Statement

The authors have declared no competing interest.

