## Supplementary Information for "Charge reversal at the Lhcb2 N-terminus impairs phosphorylation and PSI–LHCII complex formation"

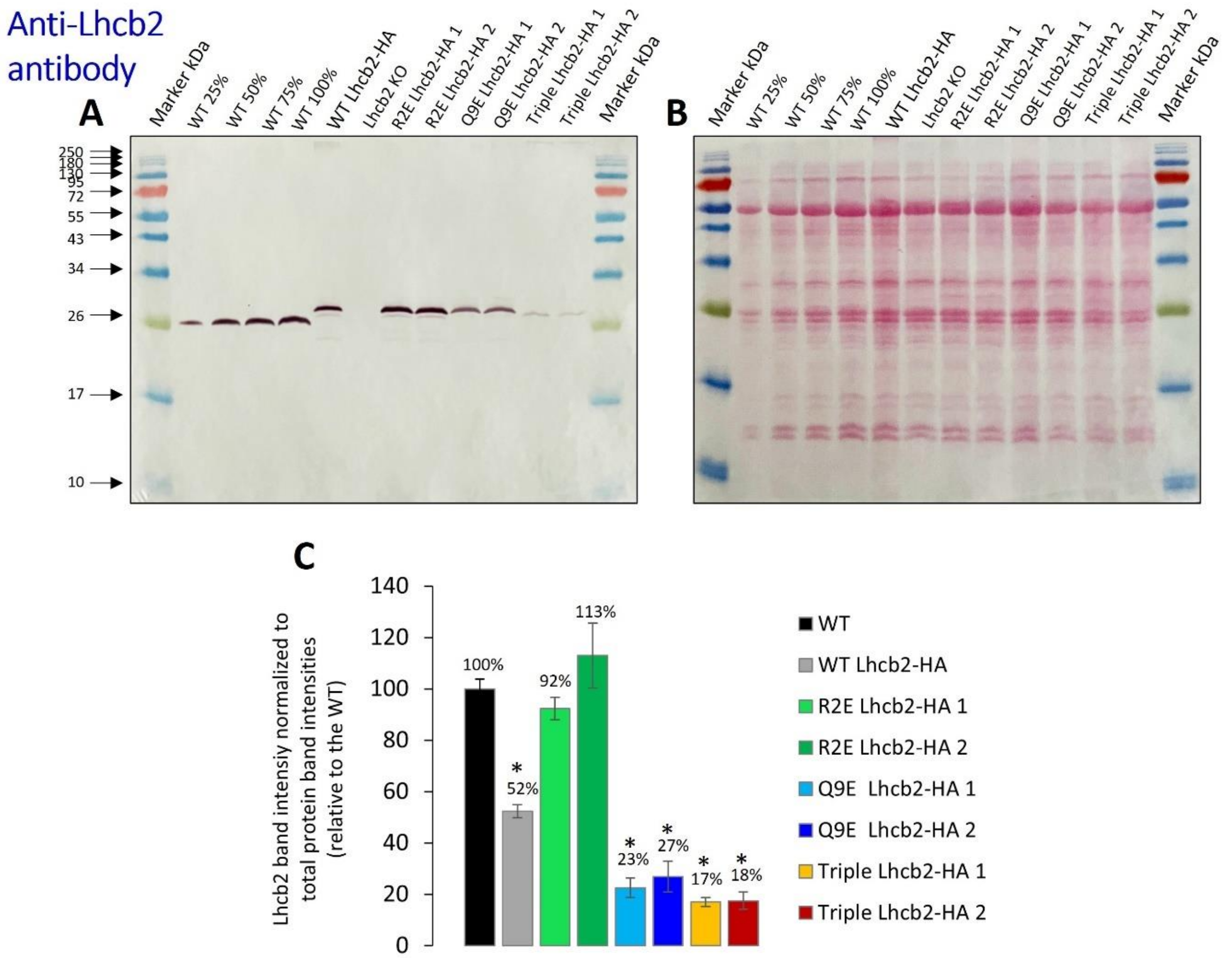


***SI Fig. S1.*** *A. Immunoblot against Lhcb2 on whole leaf extract. B. Total protein Ponceau stain of blot shown in A. C. Quantification of the immunoblot normalized to the total protein band intensity of B. Error bars represent standard error of means of four biological replicates. Statistically different values ($p < 0.05$) compared to WT are indicated with an asterisk. Statistical analysis was performed using one-way ANOVA with post-hoc Tukey test.*

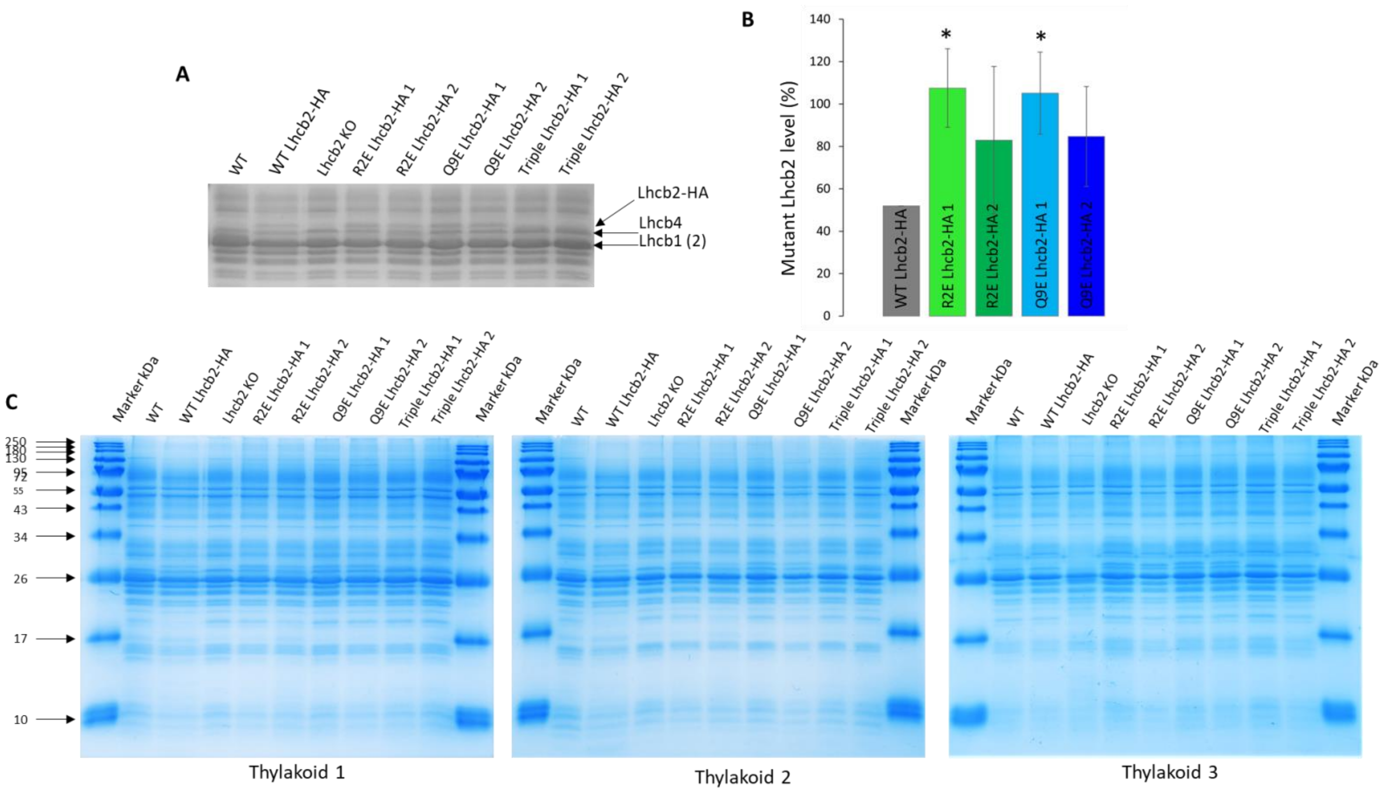


***SI Fig. S2*** *A. Coomassie stained SDS-PAGE gel of thylakoids from WT, WT Lhcb2-HA, Lhcb2 KO, R2E Lhcb2-HA (2 independent lines), Q9E Lhcb2-HA (2 independent lines), and triple mutant (D16R, E26R, E35R) Lhcb2-HA (2 independent lines). The region of the Lhc polypeptides is shown. The position of the HA-tagged Lhcb2, Lhcb4 and Lhcb1 (and Lhcb2 in the WT) are indicated. B. Lhcb2 level relative to WT plants, considering a 50% expression level for WT Lhcb2-HA, the standard error (SE) is shown based on 3 biological replicates. Statistically different values ($p < 0.05$) compared to WT-HA are indicated with single asterisk. Statistical analysis was performed using one-way ANOVA with post-hoc Tukey test. C. Entire SDS-PAGE gel of the 3 replicates.*

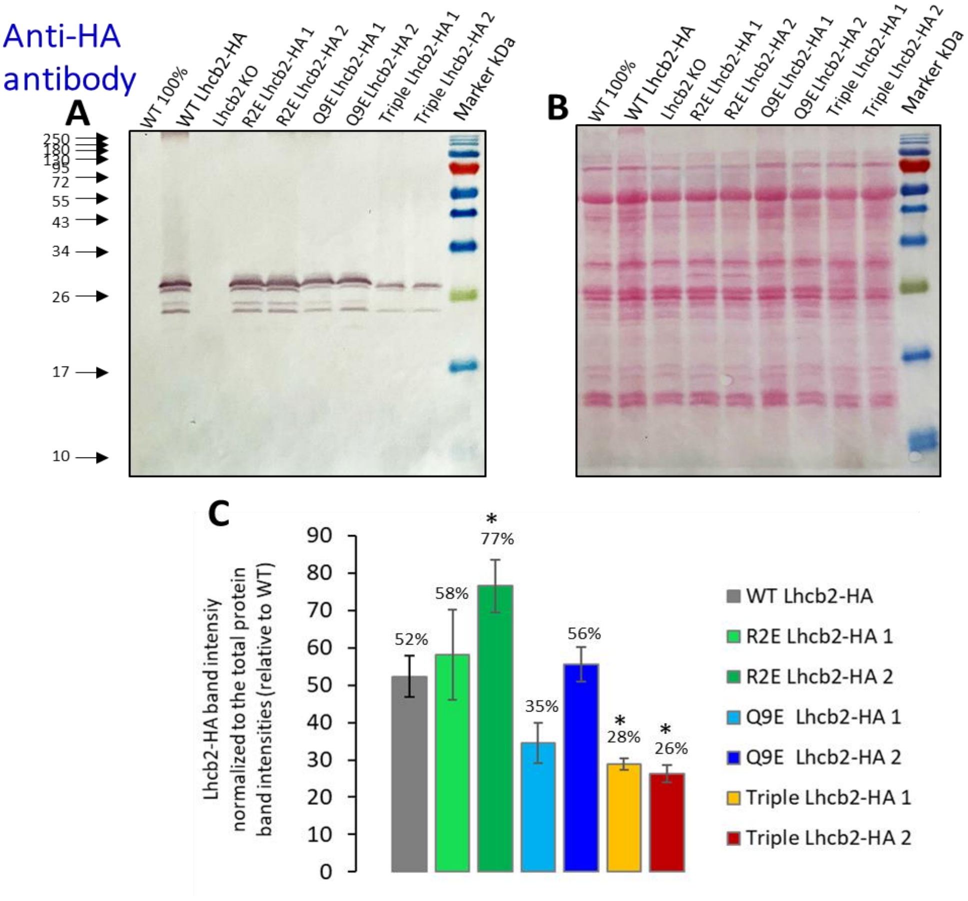


***SI Fig. S3.*** *A. Immunoblot against HA-tag on whole leaf extract. B. Total protein Ponceau stain of blot shown in A. C. Quantification of the immunoblot normalized to the total protein band intensity of B. Error bars represent standard error of means of four biological replicates. Statistically different values ($p < 0.05$) compared to WT Lhcb2-HA are indicated with an asterisk. Statistical analysis was performed using one-way ANOVA with post-hoc Tukey test.*

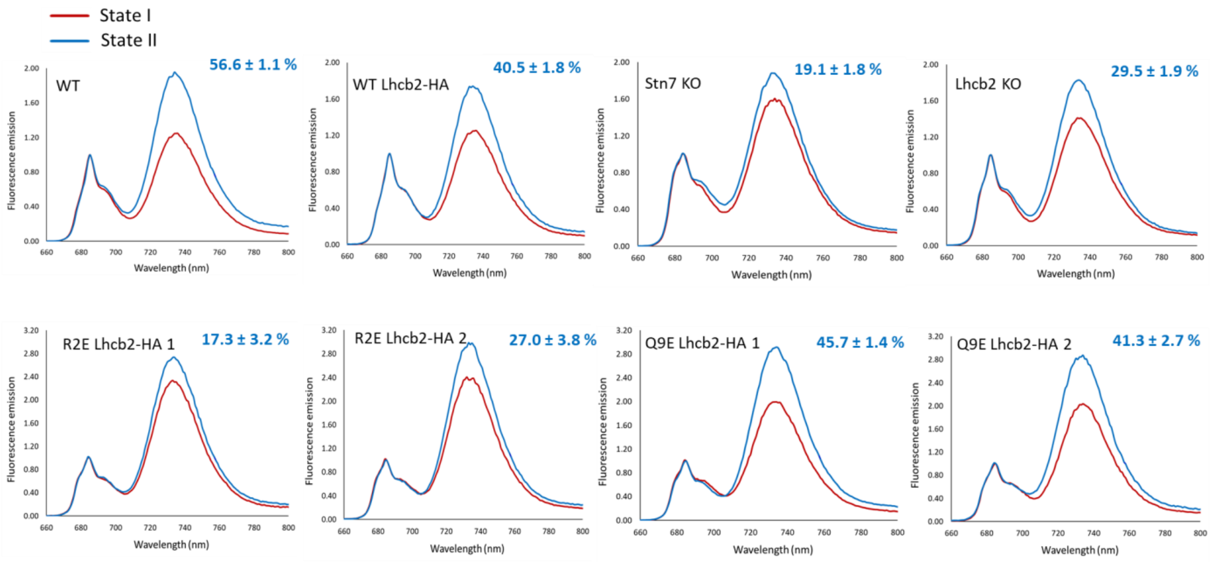


***SI Fig. S4.*** *77K fluorescence of State I and State II thylakoids. Spectra show the average of 9 technical replicates. Percentage in blue indicates the change in fluorescence at 733 nm in State II compared to State I. Spectra are normalized to the PSII emission maximum at ~685 nm.*

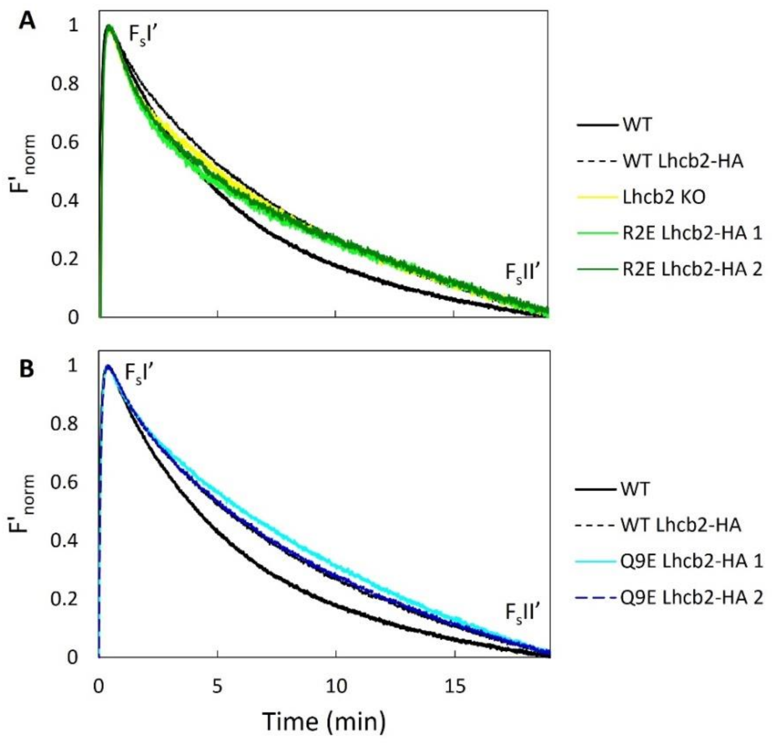


***SI Fig. S5.*** *Normalized State I to State II transition from PAM trace Figure 3 in main text. Trace at the moment when the FR light was switched off (7 to 27 minutes in Figure 3). The final value of $F_sII'$ is subtracted from the trace and next the value of FsI' is normalized to 1. A. WT, WT Lhcb2-HA, Lhcb2 KO, R2E Lhcb2-HA (two independent lines). B. WT, WT Lhcb2-HA, Q9E Lhcb2-HA (2 independent lines). An average of 5 biological replicates is shown. Note that in B the WT Lhcb2-HA and Q9E Lhcb2-HA 2 overlap.*

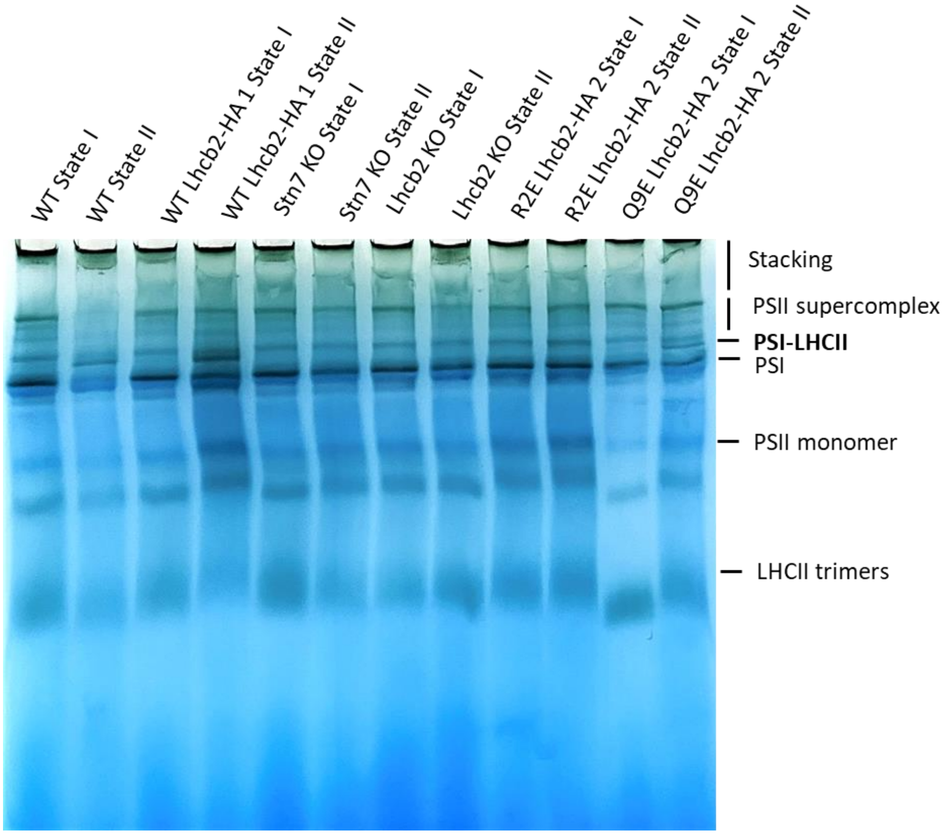


***SI Fig. S6.*** *Native PAGE as shown in Figure 4A of the main text.*

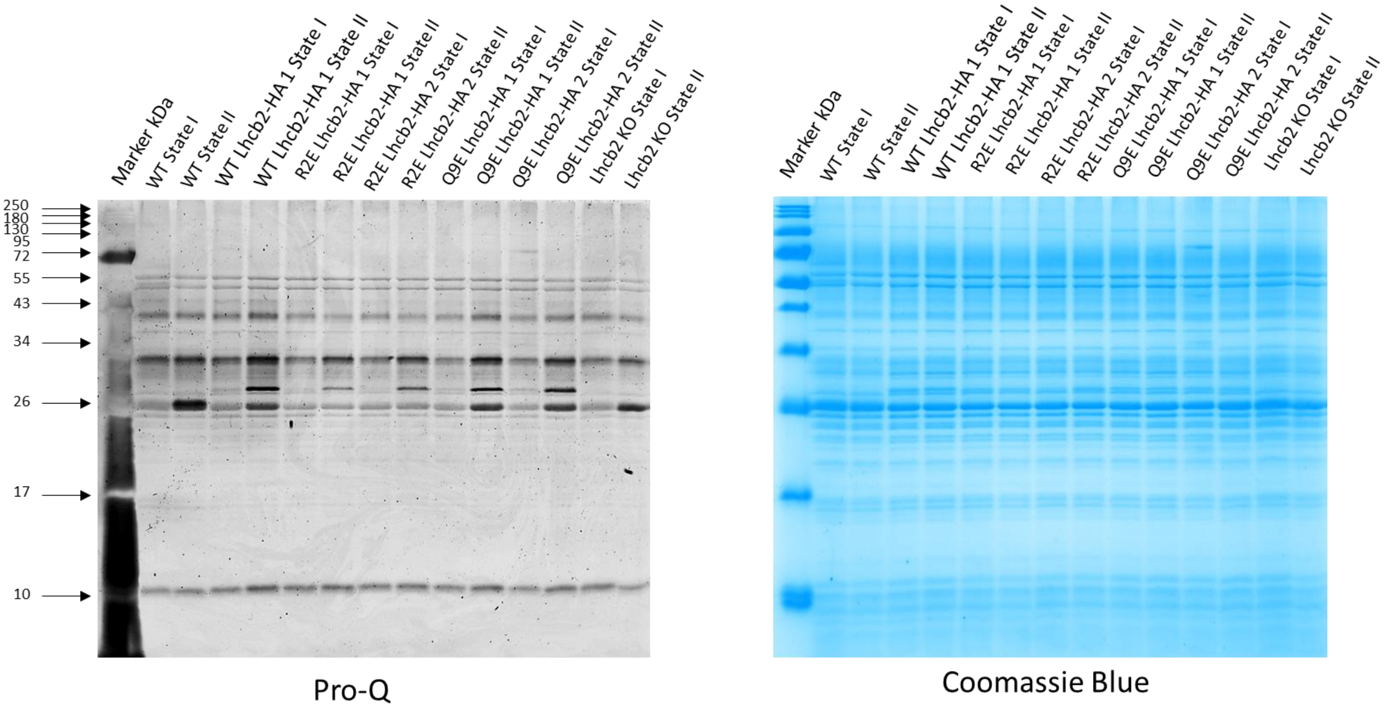


***SI Fig. S7.*** *Pro-Q Diamond phosphostain and Coomassie blue stain of SDS-PAGE gel shown in Figure 4B of main text.*

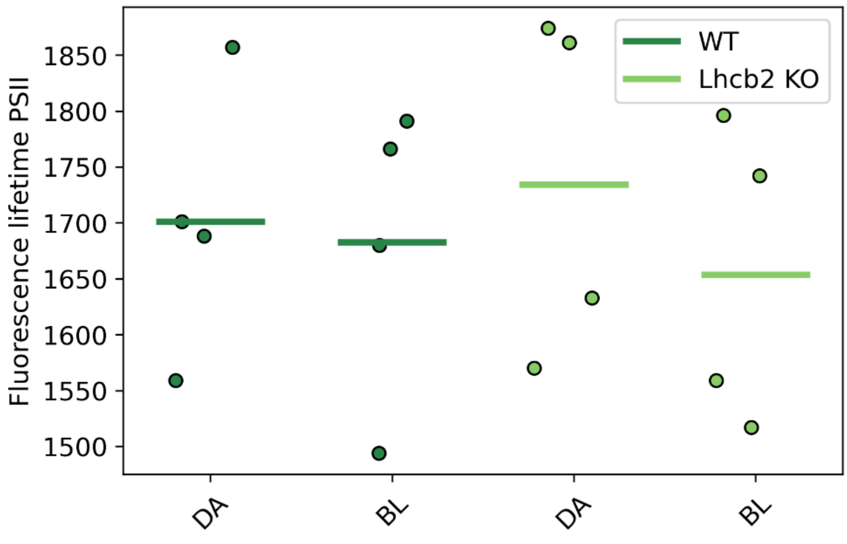


***SI Fig. S8.*** *Average PSII fluorescence lifetimes of data shown in Figure 6 of the main text.*

***SI Table S1 Analysis of Lhcb2 KO PAM data***

| Conditions | Average value ($F_m$ normalized) | Expected PSII lifetime (ns) | Expected decay rate ($ns^{-1}$) |
|---|---|---|---|
| $F_m$ | 1 | 1.8 | 0.56 |
| $F_mI'$ | 0.95 | 1.71 | 0.59 |
| $F_mII'$ | 0.92 | 1.65 | 0.61 |
| $F_sI'$ | 0.35 | 0.63 | 1.58 |
| $F_sII'$ | 0.30 | 0.54 | 1.86 |

During the State I to State II transition, the $F_S'$ value of the Lhcb2 KO mutant decreased from 0.35 to 0.30 during the illumination with red light. This decrease in $F_s'$ could be related to a shift of the excitation balance towards PSI, which relieves PSII excitation pressure, promotes re-oxidation of the plastoquinone pool, re-oxidizes the primary quinone acceptor $Q_A$, increases the fraction of open PSII reaction centers, and thereby reduces the fluorescence. Furthermore, the relocation of LHCII from PSII to PSI will decrease the PSII antenna cross-section and therefore its fluorescence. Alternatively, the decrease in $F_S'$ value can be explained by Lhcb1 phosphorylation induced fluorescence quenching. If quenching is induced in $F_S'$, then the same quencher would act during the saturation pulse. To calculate the effect of the quenching during the saturating pulse ($F_mII'$) we need to calculate the expected quenching rate that would explain the decrease in $F_S'$.

To calculate the quenching rate, we made the following assumptions:

- The contribution of PSI fluorescence is neglected, all the fluorescence is thus attributed to PSII chlorophyll *a* fluorescence.
- For chlorophyll *a* embedded in PSII the fluorescence quantum yield ($\Phi_F$) is directly related to the radiative rate ($k_r$) and the excited state lifetime ($\tau$) of chlorophyll *a* according to: $\Phi_F = \frac{k_r}{k_r+k_{IC}+k_{ISC}+k_{phot}} = k_r\tau$, with $\tau = \frac{1}{k_r+k_{IC}+k_{ISC}+k_{phot}}$.
- Under $F_m$ and $F_m'$ conditions, the rate of photochemical quenching ($k_{phot}$) is considered to be zero, while the other rates are considered not to be affected by the saturating light pulse.
- The average maximum PSII lifetime in dark adapted leaves is 1.8 ns.

- The change in PAM fluorescence values are due to changes in the fluorescence quantum yield of PSII fluorescence and not due to changes in the absorption cross-section of PSII, e.g. no relocation of the LHCII antenna during the switch from State I to State II conditions.
- The rather weak PSII quenching in $F_s$ is considered not to change the $Q_A$ oxidation status.

Based on these assumptions, the expected PSII lifetimes can be calculated from the normalized PAM values and are presented as expected lifetimes in SI Table S1. The total decay rate is given by: $k_{tot} = 1/\tau$, the expected decay rates are also presented in the table.

Under the above assumptions, the difference in the total decay rates calculated from $F_sI'$ and $F_sII'$ corresponds to a quenching rate of 1.86 – 1.58 = 0.28 $ns^{-1}$. If that identical quencher were active during the saturating pulse, the $F_m$ lifetime would be shortened from 1.71 ns to 1.15 ns, corresponding to a ≈33% reduction in $F_m$, which is substantially larger than the observed 3% reduction; therefore the observed changes in $F_s'$ are better explained by redistribution of excitation to PSI than by PSII–LHCII quenching.
